# A toolkit for studying *Varroa* genomics and transcriptomics: Preservation, extraction, and sequencing library preparation

**DOI:** 10.1101/2020.08.17.255083

**Authors:** Nonno Hasegawa, Maeva Techer, Alexander S. Mikheyev

## Abstract

**Background:** The honey bee parasite, *Varroa destructor*, is a leading cause of honey bee population declines. In addition to being an obligate ectoparasitic mite, *Varroa* carries several viruses that infect honey bees and act as the proximal cause of colony collapses. Nevertheless, until recently, studies of *Varroa* have been limited by the paucity of genomic tools. Lab- and field-based methods exploiting such methods are still nascent. This study developed a set of methods for preserving *Varroa* DNA and RNA from the field to the lab and processing them into sequencing libraries. We performed preservation experiments in which *Varroa* mites were immersed in TRIzol, RNAlater, and absolute ethanol for preservation periods up to 21 days post-treatment to assess DNA and RNA integrity.

**Results:** For both DNA and RNA, mites preserved in TRIzol and RNAlater at room temperature degraded within 10 days post-treatment. Mites preserved in ethanol at room temperature and 4°C remained intact through 21 days. *Varroa* mite DNA and RNA libraries were created and sequenced for ethanol preserved samples, 15 and 21 days post-treatment. All DNA sequences mapped to the *V. destructor* genome at above 95% on average, while RNA sequences mapped to *V. destructor*, but also sometimes to high levels of the deformed-wing virus and to various organisms.

**Conclusion:** Ethanolic preservation of field-collected mites is inexpensive and simple, and allows them to be shipped and processed successfully in the lab for a wide variety of sequencing applications. It appears to preserve RNA from both *Varroa* and at least some of the viruses it vectors.

## Background

Honey bees are one of the most economically significant agricultural resources, contributing to crop pollination, as well as providing products such as honey, propolis, and beeswax, which also contribute to local economies [1]. The agricultural department of Canada reported that in 2016, honey bee pollination contributed 2.5 billion CAD to Canadian crops [2]; thus, the decline of honey bee populations gravely threatens agricultural output.

While honey bee declines are multifactorial, they have been accelerated by the pandemic spread of the ectoparasitic mite, *Varroa destructor*, which jumped hosts from the closely related eastern honey bee (*Apis cerana*) [1, 3]. Until recently, few genomic resources for *Varroa* existed, and how it evolved post-host switch remains poorly understood [4, 5]. Furthermore, *Varroa* mites vector several viruses that impair the honey bee immune system, cause underdevelopment in honey bees, as well as cognitive impairment [6]. Viruses transmitted by *Varroa* are believed to be the primary driver of declining honey bee populations worldwide, exacerbating parasitism by *Varroa* mites themselves [1]. Many *Varroa*-associated viruses have been identified, most notably deformed-wing virus (DWV) [1, 7–9], acute bee-paralysis virus [8, 9], Kashmir bee virus [8, 9], and the black queen cell virus [8, 9]. However, much remains unknown about the biology of both *Varroa* and viruses it carries, as standard molecular methods to study them using genomic and transcriptomic tools have not been established [9].

The majority of honey bee viruses are single-stranded, positive-sense, RNA viruses. These commonly remain dormant, leaving the bees asymptomatic; thus, field diagnosis of honey bee viruses remains challenging [6, 9]. To understand viral loads of colonies, beekeepers must send samples to facilities that are equipped with proper instrumentation for diagnosis [9]. The optimal sampling method for live organisms for laboratory analysis is snap freezing and transporting on dry ice, which is frequently not possible for field workers and beekeepers [10]. Previous work exploring alternative storage conditions for honey bee RNA yielded degraded RNA in 70% ethanol, and whole honey bee RNA began to degrade within a week in RNAlater [10]. Campbell *et al*. [11] explored preservation of *Varroa* mites in RNAlater and its effect on RNA integrity. That study suggested that only when the mites are pierced does RNA remain intact, and did not explore other preservation solutions. Thus, there is limited published research regarding *Varroa* mite storage conditions and their effect on RNA and DNA integrity. Also, existing protocols for DNA and RNA extraction require pooling several mites to obtain enough material [11, 12].

We propose a new method to extract both DNA and RNA from a single mite, sufficient in both quality and quantity for downstream analysis, such as next-generation sequencing. This method allows individual analysis, which can be used to map viruses present in each individual rather than pooled samples. Standardizing a method to analyze viruses present in each *Varroa* mite will allow biogeographical mapping to help visualize and track virus trends on a global scale. Here, we explore *Varroa* mite preservation conditions in TRIzol, RNAlater, and absolute ethanol for storage periods of up to 21 days, and their effects on DNA and RNA quality by mapping to the *V. destructor* reference genome [5] and transcriptome. We propose a more affordable and feasible method for fieldworkers who have limited immediate access to laboratory facilities or equipment.

## Results

### DNA Quality and Library preparation

Extracted *Varroa* mite DNA exhibited no trend in A260/280 values among the four treatments. In general, this method introduced contaminants into the RNA, with A260/280 values among all samples ranging from 1.76 to 3.35, with a mean of 2.76. High values indicate potential contamination, though they exceeded the threshold of 1.8 that is commonly used in laboratory practice for downstream applications [13]. We inspected samples treated with TRIzol and RNAlater using DNA electrophoresis and found that they were heavily degraded 10 days post-treatment. Mean total DNA yield was 49.7 ± 58 (s.d.) ng. DNA generally remains intact for days or weeks of storage at room temperature; thus, our target for downstream analysis was the control group and ethanol groups from 21 days post-treatment to further analyze the quality by library prep, sequencing, and mapping to *V. destructor*. DNA libraries were successfully produced using a Nextera XT library preparation kit (catalog #FC-131-1096), uniquely indexed for sequencing.

### RNA Quality and Library preparation

*Varroa* mite total RNA was extracted with a mean yield of 2,157 ± 2,570 (s.d.) ng. RNA electrophoresis showed that RNA in samples preserved in ethanol remained intact even at room temperature, while RNAlater and TRIzol preserved samples had degraded heavily in both 15- and 21-day samples (Fig. 1). Control group samples, which were snap-frozen and stored at −80°C, were also well-preserved. As RNAlater fails to penetrate mites unless their exoskeletons are punctured, degradation of RNA was expected [11]. *Varroa* mites float to the surface in TRIzol, as they are less dense than the solution; thus, RNA degradation was also expected in these samples. Total RNA libraries were prepared using an NEB low-input RNA library preparation kit (catalog #E6420), uniquely indexed for sequencing

**Figure 1:**
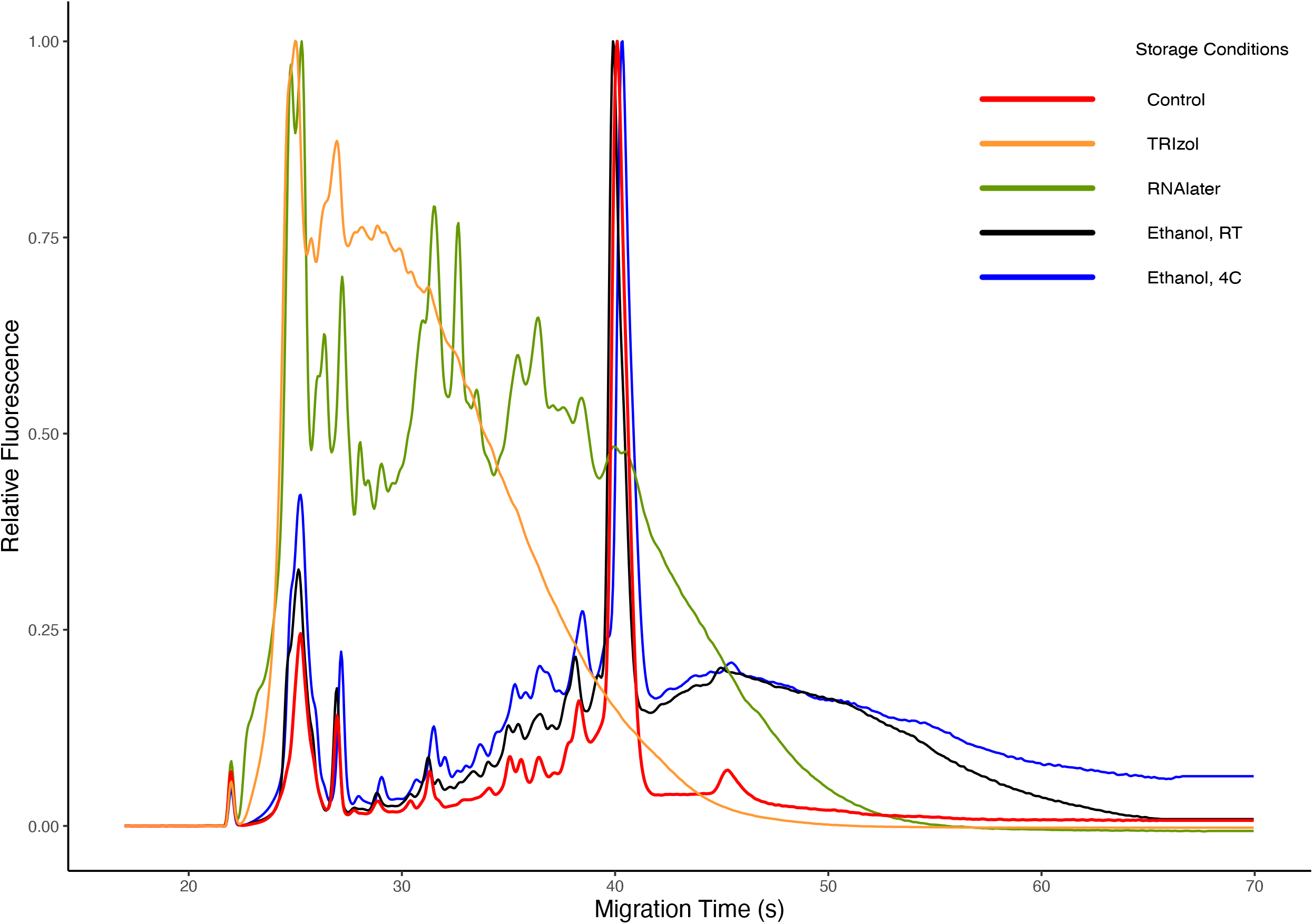
Representative bioanalyzer result of Varroa mite total RNA, extracted 21 days post-treatment. Control samples that were snap-frozen and stored at −80°C show minimal noise and a clean 18S peak, while ethanol samples at room temperature and 4°C also showed a similar 18S peak; however, with more degradation products. Mites stored in RNAlater and TRIzol had degraded and did not show a peak at 18S, indicating that RNA preservation was not successful. This suggests that weeks of storage in ethanol, even at room temperature, have little effect on RNA integrity.

### DNA and RNA Sequence Mapping

*Varroa* mite DNA libraries acquired, on average, ^~^231,000 ± 49,000 (s.d.) sequences, whereas RNA libraries averaged approximately 447,000 ± 248,000 (s.d.) sequences per library. Given the short-read length used in sequencing, inspection with FastQC revealed that there were essentially no adapters and that sequence quality was consistently high, so raw reads were not processed further. DNA libraries of mites preserved in ethanol for 15 and 21 days at room temperature and 4°C aligned to the *V. destructor* reference genome with a median of 95.49 ± 1.16 (s.d.) % (Fig. 2). Of the mapped reads, approximately 5.5 ± 0.74 (s.d.) % were PCR duplicates, an artifact of library preparation. Since ribosomal RNA depletion was not performed before library preparation, we also conducted alignments to 18S (FJ911866.1) and 28S (FJ911801.1) rRNA to ensure that ribosomal RNA (rRNA) was not abundant, possibly swamping the mRNA sample. On average, there was less than 0.001% rRNA, allowing us to proceed with general mapping to *Varroa* mite gene models. RNA sequences from *Varroa* mites stored in ethanol at both room temperature and 4°C mapped inconsistently to *V. destructor*. Mapping to the *V. destructor* reference library varied from as little as 2% to as high as 76%. We used Kraken2 [14] to classify reads to other organisms, which mapped to various species (Fig. 2). Taxon classification with Kraken2 suggested higher mapping to DWV for individuals that didn’t map well to the *V. destructor* reference library. There was no difference in the mapping rates between RNA-seq libraries prepared by our approach *vs*. those prom prior experiments that are available on the NCBI SRA database (Welch Two Sample t-test t16.4 = 0.12, p = 0.91).

**Figure 2:**
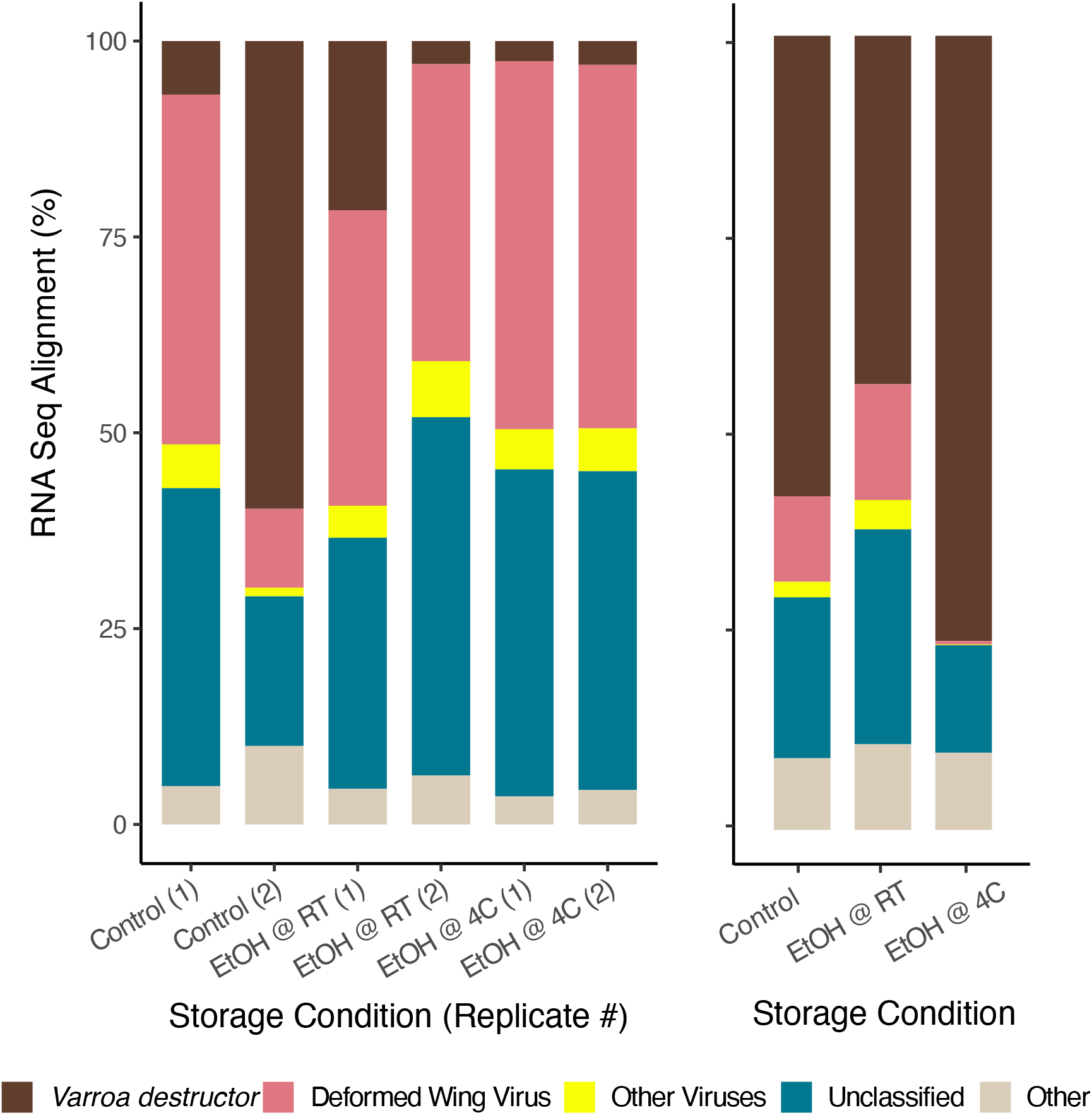
Alignment of RNA libraries from *Varroa* mite total RNA to *V. destructor* genome sequences, DWV, and other viruses. Some samples had fewer *V. destructor* reads, compared to DWV. Other viruses remained at a lower mapping percentage, and some reads did not map to any reference genome. Reads that remained unclassified may be organisms that have not yet been sequenced, library preparation artifacts, new RNA viruses for which *Varroa* mites serve as vectors, or microbes.

## Discussion

Transporting samples from the field to the lab for RNA processing often involves considerable stress and logistical complexity. To facilitate this process, we assessed effects of different preservation conditions on *Varroa* mite DNA and RNA integrity. Surprisingly, *Varroa* mite total DNA and RNA libraries mapped well to the *V. destructor* genome when bees were stored in absolute ethanol at room (Fig 2). This was a significant finding since for honey bees [10] and spiders [15], ethanol does not preserve RNA well. While butterfly [16], honey bee [10], and *Varroa* mite [11] RNA preserved well in RNAlater when specimens were ground, sliced, or pierced, intact specimens generally degraded within a few days post-treatment. Similar to these findings, we found that intact *Varroa* mite DNA and RNA stored in TRIzol and RNAlater started to degrade within 10 days, and possibly earlier.

As previously reported [17], DNA preserves well over longer periods than RNA, as long as the solution penetrates the specimen. *Varroa* mites floated to the surface in TRIzol; thus, mite DNA was not well preserved. Similarly, mites immersed in RNAlater were not well preserved because the solution does not penetrate the cuticle. Through RNA electrophoresis, we found that *Varroa* mite total RNA was completely degraded 2 weeks post-treatment (possibly earlier) when stored in TRIzol or RNAlater (Fig. 1). Surprisingly, RNA samples 21 days post-treatment immersed in ethanol at room temperature remained largely intact, allowing appropriate NGS library preparation and analysis. Ethanol RNA samples at both room temperature and 4°C mapped either to *V. destructor* or DWV, with a minimal amount of rRNA. Chen *et al*. [10] stated that whole honey bee RNA did not preserve well in RNAlater without slicing or crushing 1-week post-treatment, which is consistent with our results. Several other studies also suggest that spiders [15] and butterflies [16] preserve well in RNAlater when specimens are crushed. However, RNA integrity varies when stored in ethanol between organisms. Spiders only preserved well when crushed, and not when immersed in ethanol as whole organisms [15].

Mites must be punctured [11] if these solutions are to successfully preserve sample RNA, as for honey bees [10]. However, it is not practical for beekeepers to puncture each specimen, particularly in the field, and TRIzol and RNAlater are only available through vendors in laboratory reagents and are much more expensive than ethanol. For these reasons, we recommend the use of ethanol, which is readily available from chemical suppliers. *Varroa* mites yielded high-quality DNA and RNA for NGS analysis for weeks when stored in ethanol at room temperature or at 4°C, which should be sufficient for collection and shipping. Though we terminated the experiment at three weeks, it seems likely that both DNA and RNA are stable for much longer, given the minimal degradation we observed at 21 days.

Although RNA libraries mapped well to *V. destructor* and DWV, there was still a portion of RNA sequence that was not classified. These unclassified sequences may contain new, unidentified RNA viruses that the *Varroa* mites are vectoring. They may also belong to organisms that have not been sequenced or may represent library preparation artifacts. Though it is beyond the scope of our current study, and not entirely feasible with 50-bp reads used in these experiments, exploring the unclassified region of RNA sequences will be beneficial in understanding the complicated host-parasite relationship of honey bees and *Varroa* mites, opening new avenues to honey bee health.

There are several limitations we must take into account. Most importantly, our sample size is limited to n=6 per treatment per time point. However, the number of samples we processed for library preparation and sequencing resulted in high-quality mapping to the reference genome, and none of the samples deviated strongly from the rest, suggesting we have a good representation of the treatments. Quality of ethanol must also be considered. In laboratory experiments, we require high-grade alcohol for quality control of samples. So, while we used high-grade absolute ethanol in this study, this quality of ethanol is not readily available worldwide. We suspect that lower quality ethanol, particularly if contaminated with impurities such as methanol, may harm preservation as they cause more hydration and further denature nucleic acids [18].

Although both DNA and RNA integrity were retained at room temperature when immersed in ethanol, it is more realistic and practical to store and ship samples within 2 weeks (15 days post-treatment). We advise beekeepers and field workers to store mite samples in the refrigerator when possible, and at room temperature when this is not possible. Samples may also be shipped without refrigeration, reducing shipping costs. Additionally, as we observed 21 days of stable storage, shipping delays should be considered if RNA integrity must be maintained for longer periods for high-quality NGS and other downstream analyses. In short, we found that a possible alternative to snap freezing *Varroa* mites and storing at −80°C is immersion in absolute ethanol for up to 21 days. By immersing in absolute ethanol, *Varroa* mite DNA and RNA are well preserved at both room temperature and at 4°C, which allow for more flexible sampling and storage conditions. We believe that methods presented in this study will lead to insights in *Varroa* genomics and population biology and will facilitate studies of the viruses vectored by *Varroa*.

## Conclusion

We found that *Varroa* mite DNA and RNA were adequately preserved in absolute ethanol for up to 21 days, and produced high quality DNA and RNA libraries when sequenced, mapping to the *Varroa* genome and to other taxa. On the contrary, when *Varroa* mites were preserved in TRIzol and RNAlater, the mite DNA and RNA degraded within the first 10 days, possibly earlier, likely as a result of poor penetration through the exoskeleton. Ethanolic preservation of Varroa mites is inexpensive and uses a readily available reagent, thus allowing specimens to be shipped and processed for a wide variety of sequencing applications. In addition, ethanol also preserves viruses that Varroa vectors, most notably DWV. We propose that ethanolic preservation can replace cryopreservation, providing a more tractable method for preserving DNA and RNA quality.

## Methods

### Mite Collection

In March 2020, *Varroa* mites were collected from managed hives in Onna village, Okinawa by removing honey bees from 2 frames onto a tray with icing sugar, and shaking the tray to remove mites from honey bees (Fig. 3). From the site of collection to the laboratory, mites were in icing sugar. Once at the laboratory, we discarded dead mites. Mites were divided into the following treatment preservation conditions: 1) snap-frozen and stored at −80°C 2), immersed in absolute ethanol at room temperature, 3) immersed in absolute ethanol at 4°C, 4) immersed in TRIzol (Thermo Fisher Scientific, Tokyo, Japan) at room temperature, and 5) immersed in RNAlater (Thermo Fisher Scientific, Tokyo, Japan) at room temperature. When immersed in a solution, 500 μL was used and each treatment consisted of 6 mites placed in separate tubes. Each treatment group was subjected to DNA and RNA extractions at intervals of 5, 10, 15, and 21 days for subsequent DNA and RNA quantity and quality evaluations. Snap-freezing and storage at −80°C were chosen for the control group as it is a widely used method for specimen preservation [6, 15, 16]

**Figure 3:**
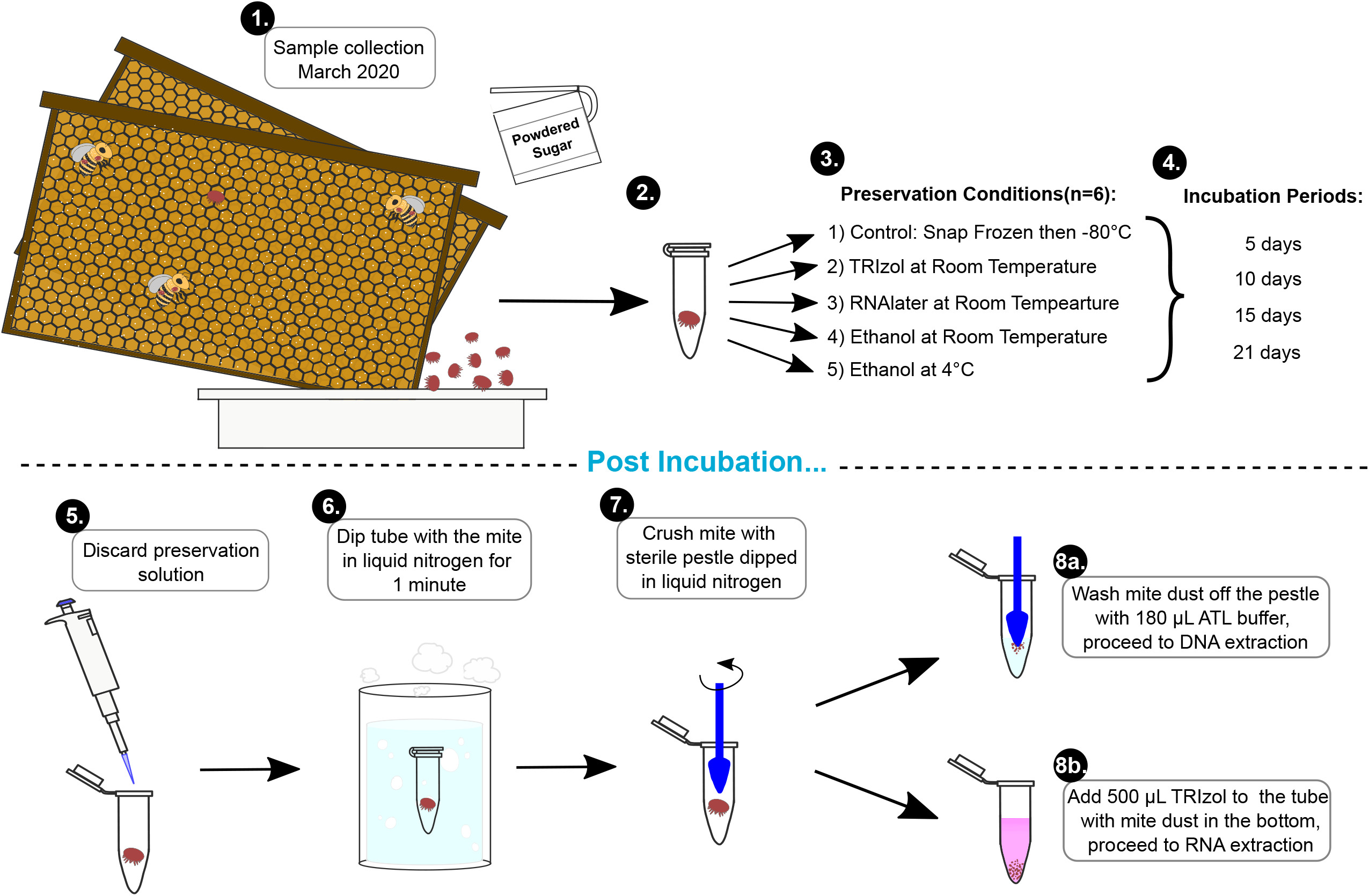
Generalized experimental workflow. 1) Varroa mites were collected from an apiary located at the Onna village office in Okinawa, Japan, March, 2020. Mites were shaken off the honey bees by sprinkling powdered sugar on the frame. Bees were collected into a tray and shaken to remove mites. The tray containing powdered sugar and Varroa mites was transported back to the laboratory, within 30 minutes, discarding dead mites on arrival. 2) Individual mites were placed in separate 1.5-mL microcentrifuge tubes and 3) were immersed in 500 *μ* L of a preservative solution or snap-frozen and stored at −80 ° C until processing. 4) Incubation periods were 5, 10, 15, and 21 days in respective preservation methods. 5) Preservation solution was discarded after the incubation period, and 6) samples were immersed in liquid nitrogen for a minute, then 7) crushed with a sterile pestle that was also immersed in liquid nitrogen. Each sample was split in two tubes, 8a) mite dust on the pestle was washed with ATL buffer into a new tube and used for DNA extraction with a QIAamp DNA extraction kit and 8b) TRIzol was added to the tube containing mite dust and subjected to RNA extraction using TRIzol.

### Mite Preparation for Extractions

Mites were removed from the solution in which they were stored to new 1.5-mL tubes (Eppendorf catalog #0020125215) and chilled in liquid nitrogen. A clean, autoclaved pestle (Sigma Aldrich catalog #Z359947) was also chilled in liquid nitrogen to grind each mite in its tube. By visual inspection, ground mite powder that remained on the pestle was used for DNA extraction, ensuring that about half of the mite homogenates remain in the tubes for RNA extraction (Fig. 3).

### DNA Extraction

Mite DNA was extracted using a QIAamp DNA Micro kit (Qiagen, Tokyo, Japan), according to the manufacturer’s protocol. DNA quantity was determined using a Qubit fluorometer (Thermo Fisher Scientific, Tokyo, Japan), and quality was evaluated by the A260 / A280 ratio using a NanoDrop spectrophotometer (Thermo Fisher Scientific, Tokyo, Japan). Eluted DNA was stored at −20°C until further applications.

### RNA Extraction

Mite RNA was extracted using TRIzol according to the manufacturer’s protocol. Due to the small amount of mite present, the protocol was modified by using 50% of specified reagent volumes. Total RNA quality and quantity were evaluated using absorbance ratios of A230/260 and A230/280 on a Nanodrop spectrophotometer.

### Library Preparation

#### DNA Library Preparation

DNA libraries were prepared using a Nextera XT DNA Library Prep kit (Illumina, Tokyo, Japan) according to the manufacturer’s protocol, with optimization for mite DNA, using reagents at 20% of their specified volume. DNA was visualized by running electrophoresis on 1% agarose gels, for 20 min at 135 V

#### RNA Library Preparation

RNA libraries were prepared using an NEBNext Single Cell/ Low-Input RNA Library kit (New England BioLabs, Tokyo, Japan), according to the manufacturer’s protocol. Purified cDNA was indexed with i5 and i7 primers (catalog #7600S, New England BioLabs, Tokyo, Japan), and then purified and size selected for a range of 400 to 2000 bp using 11% and 11.5% PEG and DynaBeads [19].

Both DNA and RNA libraries, as well as total RNA, were analyzed with a Bioanalyzer 2100 (Agilent, Tokyo, Japan) or a 4200 Tapestation (Agilent, Tokyo, Japan). For Bioanalyzer, high-sensitivity DNA kits and RNA pico kits (Agilent, Tokyo, Japan) were used, while for the Tapestation, a high-sensitivity D5000 kit (Agilent, Tokyo, Japan) was used.

### Sequencing and Analysis

A MiSeq (Illumina, Tokyo, Japan) was used to perform both DNA and RNA sequencing, at 50 cycles and read with single read-only. We were interested in validating the protocol and estimating mapping percentages, so longer read lengths or higher coverage were not necessary. Raw sequence data from the MiSeq were first analyzed with FastQC [20] for quick quality control to see if adapter removal was necessary and to ensure that sequenced data was of sufficient quality. DNA sequence data were then analyzed using Bowtie2 [21], Samtools [22], and Picard tools [23] to map to the reference genome (Vdes_3.0 [5]) and to identify duplicates. RNA sequence data were analyzed using Bowtie2 and Samtools, and then taxonomically classified using Kraken [14]. We compared this protocol with previously published RNA-seq libraries deposited on NCBI, which were mapped to the reference genome in a similar way (SRR8867385 [24], SRR5109825 & SRR5109827 [8], SRR533974 [25], SRR5760830 & SRR5760851 [26], SRR8100122 & SRR8100123, SRR5377267 & SRR5337268, and SRR8864012 [27]).

## Abbreviations

DWV: deformed wing virus

## Declarations

### Ethics approval and consent to participate

Not applicable. Permissions for sample collections were unnecessary as they are managed on site by colleagues.

### Consent for publication

All authors have consented to publishing this work.

### Author contributions

All authors designed the study. NH and MT performed the laboratory work. NH performed the analysis and wrote the first manuscript draft, to which all authors contributed.

## Acknowledgements

We are grateful to Miyuki Suenaga for help in the lab, the sequencing centre at OIST, and to the Honey & Coral project of the Onna Village for help in maintaining the bee colonies. We are grateful to Alexandra Sébastien for conducting pilot experiments that helped guide this project. We are grateful to Steven D. Aird for editing the manuscript.

## Data availability

Raw sequence data are available under DDJB/NCBI BioProject PRJDB10252. To compare our results using the following sequences obtained from sequence read archive NCBI:

SRR8867385 [24]: RNA-seq of pooled *V. destructor* adults https://www.ncbi.nlm.nih.gov/sra/?term=SRR8867385 SRR5109825 & SRR5109827 [8]: Small RNA of *Varroa destructor*: South Africa https://www.ncbi.nlm.nih.gov/sra/?term=SRR5109825
SRR533974 [25]: RNA-seq of *Varroa destructor* https://www.ncbi.nlm.nih.gov/sra/?term=SRR533974
SRR5760830 & SRR5760851 [26]: GSM2686390: R245-9; Varroa destructor; RNA-Seq https://www.ncbi.nlm.nih.gov/sra/?term=SRR5760830
SRR8100122 & SRR8100123: RNA-seq of *Varroa destructor* https://www.ncbi.nlm.nih.gov/sra/?term=SRR8100122
SRR5377267 & SRR5337268: VD_FemaleFound1 https://www.ncbi.nlm.nih.gov/sra/?term=SRR5377267
SRR8864012 [27]: RNA-seq of *Varroa destructor* adult female https://www.ncbi.nlm.nih.gov/sra/?term=SRR8864012

